# Large-scale synthetic data enable digital twins of human excitable cells

**DOI:** 10.1101/2025.09.03.674034

**Authors:** Pei-Chi Yang, Mao-Tsuen Jeng, Deborah K. Lieu, Regan L. Smithers, Gonzalo Hernandez-Hernandez, L. Fernando Santana, Colleen E. Clancy

## Abstract

Individual variability shapes how diseases manifest, how patients respond to therapy, and how rare phenotypes arise. Conventional experimental approaches obscure variation by averaging, which limits mechanistic insight and predictive accuracy. We present a computational framework that builds digital twins of human-induced pluripotent stem cell-derived cardiomyocytes from a single optimized voltage clamp experiment. The framework depends on massive synthetic datasets comprising simulated cells that span broad ionic and electrophysiological ranges. These synthetic data make it possible to control parameters precisely, explore biological variability comprehensively, and train models beyond the limits of experimental data. A neural network trained on synthetic data then inferred biophysical parameters from experimental recordings from live cells, reproducing distinct electrophysiological features. Our study unites computational modeling, data simulation, and learning to enable scalable, precise, individualized cardiac electrophysiology modeling and can be readily extended to any electrically active cell type.

## Introduction

Individual variability determines the emergence and severity of disease, as well as individual therapeutic response in the context of both inherited and acquired conditions^1–3^. Traditional preclinical models are designed to minimize variability to isolate specific biological effects. While this reductionist approach simplifies experimental interpretation, it fails to reflect the heterogeneity that exists in human populations. The consequence is a persistent translational gap, an incomplete mechanistic understanding of disease, and a higher risk of adverse drug reactions^4–8^. Digital twins constitute a mechanistic approach to integrate individualized data with simulation-based inference to predict physiology and therapeutic outcomes at the level of an individual cell^9–12^.

Human induced pluripotent stem cell-derived cardiomyocytes (iPSC-CMs) are a powerful platform for studying electrophysiology in a patient-specific context^13–19^. However, experimental measurements of iPSC-CM action potentials (APs) or ionic currents provide only partial information about the underlying mechanisms^20–24^. The relationship between measured signals and the large number of biophysical parameters governing ion channel kinetics, calcium handling, and membrane transport is complex and nonlinear, making direct parameter estimation from small experimental datasets impossible. Existing fitting approaches are slow, require extensive specialized expertise, and are not scalable to capture individual variability^25–31^.

Mathematical models of cardiac electrophysiology comprise a complementary framework to link cellular dynamics and underlying ionic mechanisms^32,33^. Personalized digital twins have not yet been realized in part because of ongoing challenges to parameterize models accurately and quickly from cell-specific experimental data. Recent advances in deep learning now make it possible to solve the associated complex inverse problem by deriving the mappings from high-dimensional time-dependent recordings to the underlying biophysical parameters. The key is the use of massive synthetic datasets from large *in silico* cell populations to generate the training data necessary for accurate, generalizable inference. It is not possible to develop a comparable process with purely experimental data.

Here, we present an integrated computational modeling and simulation, deep learning framework, and experimental recordings demonstrating the generation of high-fidelity, electrophysiology-based digital twins of iPSC-CMs from a single optimized voltage-clamp protocol. Large-scale synthetic model iPSC-CM populations were used to train a fully connected neural network to predict 52 biophysical parameters governing six major ionic currents. A deep learning-guided optimization loop was used to design voltage-clamp protocols that maximize information content for parameter recovery. The primary goal of this framework is to enable population-level interpretation of electrophysiological variability by mapping experimental recordings to underlying ionic parameters. The models reconstructed from the inferred parameters reproduce the AP waveform, calcium dynamics, and the contributions of individual ionic currents. When applied to experimental data, this approach generates high-fidelity digital twins of experimental recordings with high accuracy across temperatures and morphological variability.

This work addresses a major translational challenge by providing a scalable, temperature-agnostic, morphology-inclusive method for personalized cardiac modeling to bridge the gap between experimental recordings and mechanistic insight. We present a framework for predictive digital twin applications in precision cardiovascular medicine and beyond, since the method can be applied to any electrically active cell type. By uniting protocol optimization, large-scale synthetic training, and deep learning-based inference, this work advances the field toward scalable digital twin technology with broad potential for basic research, preclinical testing, and personalized cardiovascular medicine.

## Results

### Variability and Digital Twin Modeling of iPSC-CMs

To assess the electrophysiological variability inherent in populations of iPSC-CMs, we first generated a simulated population of 200,000 spontaneously beating cells by applying ±40% random variation to 52 biophysical model parameters (see Supplementary Information). These parameters govern six key ionic currents central to cardiac AP generation and repolarization: I_Kr_, I_CaL_, I_Na_, I_Ks,_ I_K1,_ and I_f_. Examples of population variability are shown in 20 APs in **Figure 1A**. With a broad range of parameter perturbations, extensive variability was observed in the model population, as seen in experimental iPSC-CM preparations.

**Figure 1:**
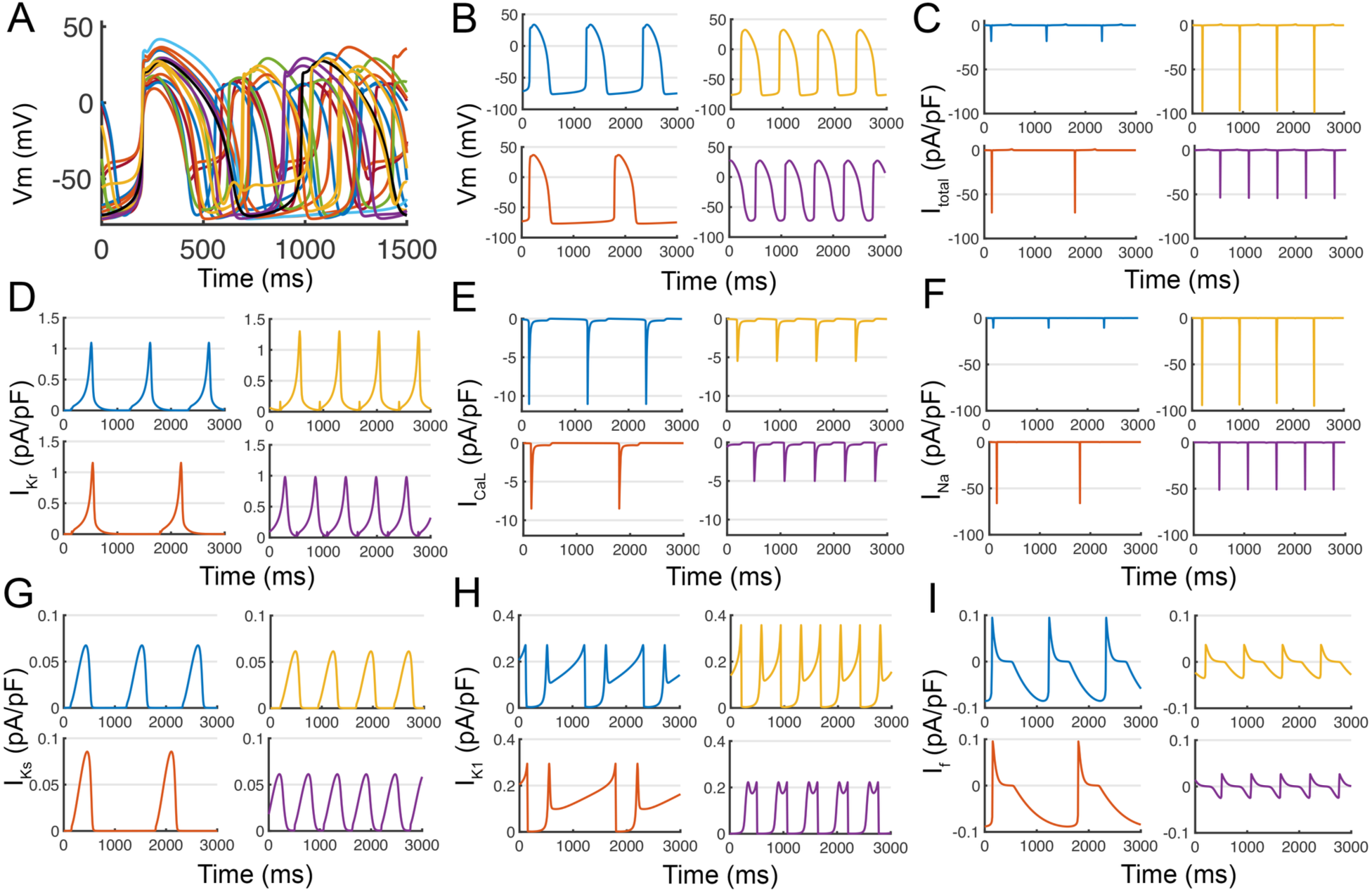
Huge variability captured in simulated iPSC-derived cardiomyocyte populations. **(A)** To illustrate population variability, 20 action potentials (APs) were shown, each resulting from ±40% random variation applied to 52 parameters governing six key ionic currents (I_Kr_, I_CaL_, I_Na_, I_Ks,_ I_K1,_ and I_f_) in the baseline Kernik iPSC-CM model ^44^, within a simulated population of 200,000 spontaneously beating cells. **(B)** These perturbations yielded a wide spectrum of APs, with substantial variation in both waveform and frequency. The corresponding total ionic current **(C)** and its decomposition into individual current components (**D–I**) are shown.

The simulations revealed a wide distribution of AP morphologies and spontaneous firing frequencies (examples in **Figure 1B**). APs varied substantially in upstroke velocity, plateau duration, and repolarization slope, with some cells exhibiting prolonged plateaus while others demonstrated rapid repolarization and shorter cycle lengths. Differences in spontaneous beat rate reflected variability in diastolic depolarization rates, which were closely linked to variations in I_f_ amplitude and I_K1_ strength across the simulated population.

To dissect the mechanistic drivers of electrophysiological diversity, we examined the total membrane current (I_Total_) shown in Figure 1C and decomposed it into individual ionic components for representative cells across the population (**Figure 1D–I**). The heterogeneity in total current waveforms was accompanied by diverse current profiles across I_Kr_, I_CaL_, I_Na_, I_Ks,_ I_K1,_ and I_f._ Some cells exhibited larger I_CaL_ amplitudes and prolonged calcium entry during the plateau, while others showed enhanced I_K1-_ or I_Kr_-mediated repolarizing currents. Differences in I_Na_ upstroke magnitude and I_f_ driven diastolic depolarization contributed to variation in excitability and cycle length. Together, these simulations underscore the high degree of functional variability that can emerge from relatively modest variation in channel kinetics and conductance properties. The synthetic cell population mirrors the experimental heterogeneity observed in iPSC-CM populations and underscores the likely importance of accounting for cellular variability in predictive modeling and ultimately, in individual drug response evaluation.

We next developed a deep learning-based workflow to generate fully parameterized models of iPSC-CM electrophysiology from whole-cell current data obtained with one simple voltage-clamp protocol (**Figure 2**). To create the training dataset, we generated a large synthetic iPSC-CMs population by introducing random variations to 52 biophysical parameters governing six key ionic currents in the baseline model: I_Kr_, I_CaL_, I_Na_, I_Ks,_ I_K1,_ and I_f_. The parameter perturbations produced an array of simulated APs and whole-cell currents, reflecting broad physiological variability (**left**).

**Figure 2.**
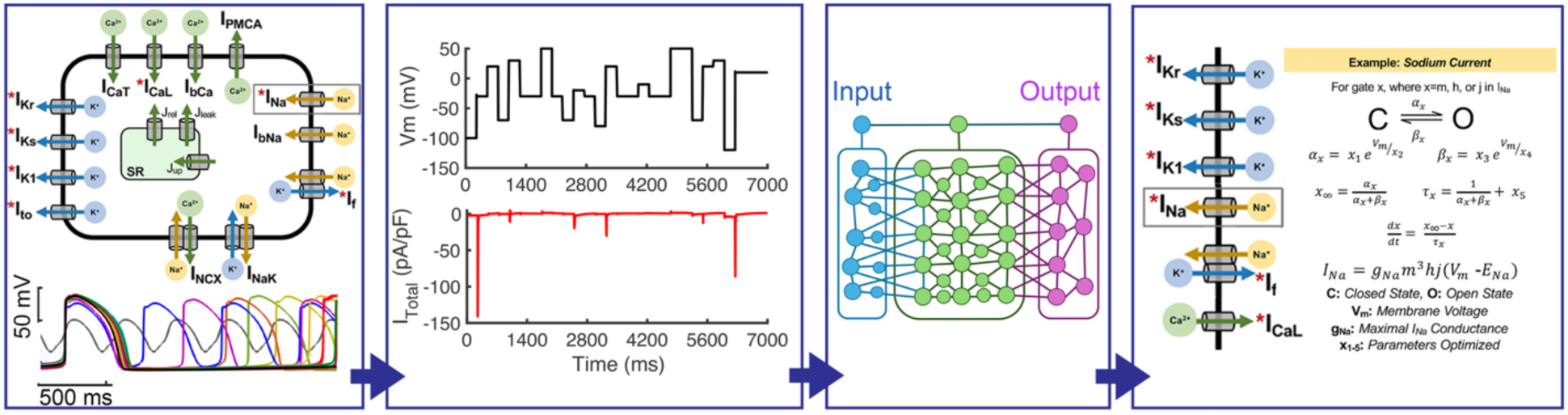
Digital twins of human iPSC-CMs from one simple voltage-clamp recording. A large population of synthetic iPSC-CMs (left) is generated by introducing variation to 52 biophysical parameters governing key ionic currents in the baseline model: I_Kr_, I_CaL_, I_Na_, I_Ks,_ I_K1,_ and I_f_ (*highlighted with red asterisks* in the schematic on the right). A computationally optimized voltage-clamp protocol (black trace, left middle) is applied to generate a distinct whole-cell current (I_Total,_ red trace), enabling cell-wide excitation of key ion channels. The simulated whole-cell currents I_Total_ from large synthetic datasets serve as inputs to a fully connected neural network trained to map raw current responses to the 52 underlying model parameters. The deep learning model is trained to predict optimized parameter values by inferring gating kinetics and maximal conductance for each ionic species. An example formulation for the fast sodium current (I_Na_) is shown, with inferred parameters (x₁–x₅) contributing to gating and conductance (right).

As part of the workflow, a voltage-clamp protocol (black trace in left middle) was applied to each synthetic cell to generate complex current responses to dynamic membrane potential input. The protocol was designed to sequentially engage major ionic currents in the synthetic cell, ensuring that the training data contained informative current signatures for each underlying conductance. The resulting total current traces (I_Total_) served as inputs to a fully connected neural network (red trace in **left middle)**.

The network was trained to map the raw current inputs to the 52 underlying model parameters, with training performance evaluated by mean squared error (MSE) on held-out test datasets. An example parameterization for the fast sodium current (I_Na_) illustrates that the network infers gating kinetic parameters and maximal conductance (**right**).

### Deep Learning Approaches for Voltage Clamp Optimization and Parameter Estimation

To maximize the information content of experimental recordings for parameter inference, we developed a deep learning-guided strategy to design a computationally optimized voltage-clamp protocol (**Figure 3**). Starting from a broad set of candidate voltage steps, the algorithm iteratively evaluated the MSE between predicted and true parameters across large synthetic iPSC-CM populations. In each cycle, the most informative test potential was selected and incorporated into the evolving protocol, while less informative steps were discarded.

**Figure 3:**
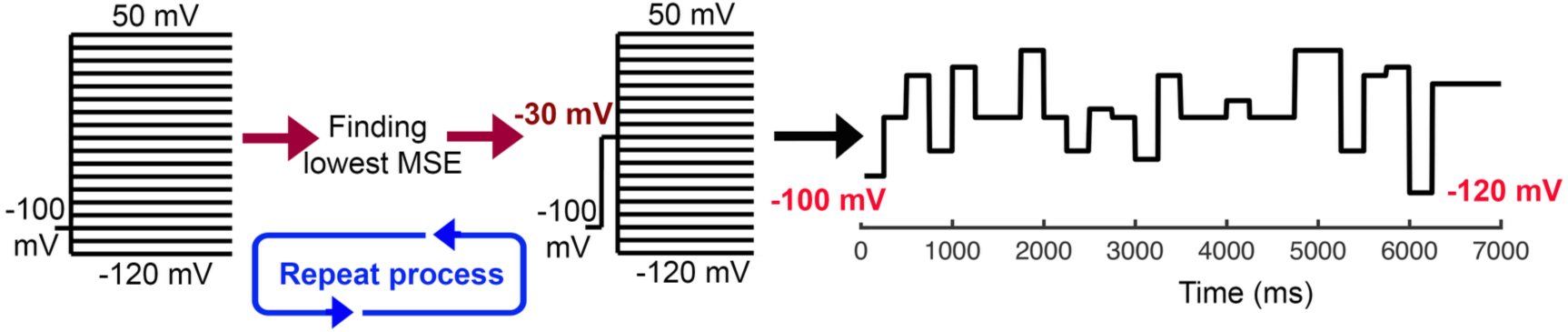
Deep learning guided optimization of a voltage clamp protocol. Each iteration began with a −100 mV holding potential for 250 ms, followed by sequential testing potentials from −120 mV to +50 mV in 10 mV increments. A total of 200,000 synthetic samples were generated per training cycle. The optimal testing potential, identified by the lowest MSE from the deep learning model, was then applied for 250 ms. This optimization cycle repeated every 7000 ms. At 6000 ms, the potential was transiently stepped to −120 mV for 250 ms before resuming the next testing potential sequence. The schematic illustrates the iterative loop of model evaluation, MSE-based selection, and protocol updating, leading to the optimized composite voltage.

Each cycle began with a -100 mV holding potential for 250 ms, followed by systematic sweeps of test potentials from -120 mV to +50 mV in 10 mV increments. A population of 200,000 computational iPSC-CMs with randomized parameter sets was simulated under each protocol iteration, and the resulting whole-cell currents were used to train the deep learning model to predict all 52 ionic parameters. The potential producing the lowest MSE was then selected as the optimal step and applied for 250 ms before the sequence resumed. This iterative optimization loop was repeated every 7000 ms, with a transient step to −120 mV at 6000 ms to probe hyperpolarization-activated channels.

This adaptive process converged on a composite voltage-clamp waveform that effectively exposed the amplitude and kinetics of all six major ionic currents while substantially reducing voltage variability through the protocol duration. The optimized sequence preserved temporally distinct current signatures for I_Kr_, I_CaL_, I_Na_, I_Ks,_ I_K1,_ and I_f,_ enabling robust parameter estimation.

When applied to held-out synthetic iPSC-CMs test data, the optimized voltage clamp protocol (**Figure 4A, left)** produced digital whole-cell currents (**Figure 4A, right)** that were effective as network training data, confirming that the protocol provided sufficient information for accurate model personalization. Training data size was still influential and determined predictive accuracy, with improvements in trained networks across datasets containing 1,000, 10,000, and 200,000 synthetic cells (**Figure 4B**). For the smallest dataset (1,000 cells), the test error plateaued at an MSE of 0.0036, with a median mean absolute error (MAE) of 0.041 across parameters. The test error curve exhibited an asymmetric U-shape across epochs, suggesting overfitting to the limited training set (**4B, left**). Increasing the dataset size to 10,000 cells improved test performance, reducing the median MAE to 0.038 (a 7.3% decrease) and narrowing the error distribution (**4B, middle**). The largest dataset of 200,000 cells had very low prediction errors, with training and test curves converging to a median mean absolute error (MAE) of 0.02 (**4B, right**). Again, the optimized protocol resulted in corresponding MAE distributions that narrowed substantially with increasing dataset size, indicating improved stability and generalizability of parameter predictions.

**Figure 4.**
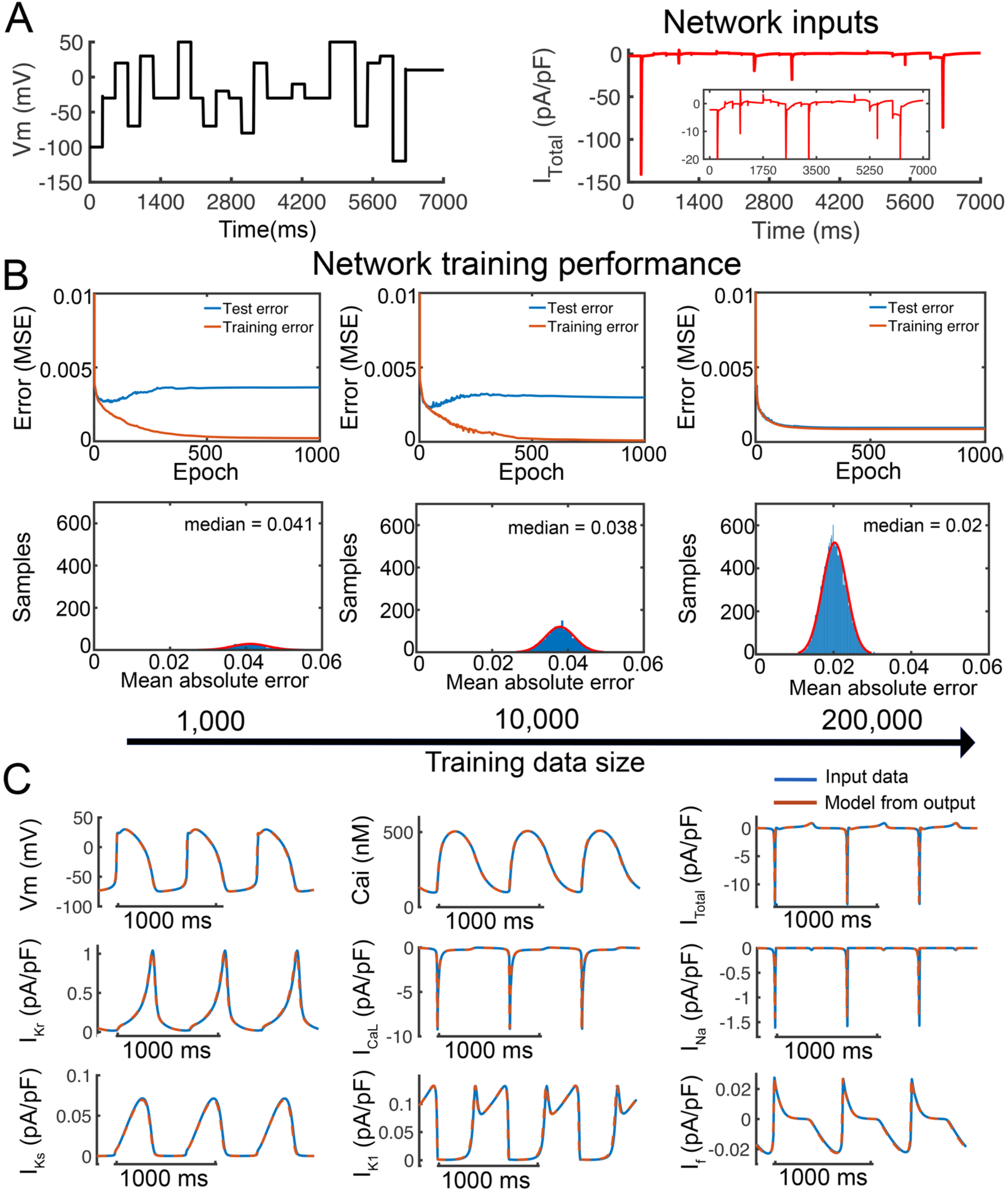
Massive synthetic data to train and test a deep learning algorithm for ion channel parameter estimation. **(A)** A deep learning-derived optimized voltage-clamp protocol (left) was designed to activate a broad set of ionic currents in iPSC-CMs by applying dynamic membrane potential steps (black trace), producing distinctive whole-cell current responses (I_Total_, red trace). The inset highlights fine-scale current kinetics captured during the protocol. These I_Total_ serve as network inputs for parameter inference. **(B)** Prediction accuracy for individual model parameters was evaluated using training datasets of 1,000, 10,000, and 200,000 synthetic cells (left to right), generated with a ±20% parameter perturbation range. For the smallest dataset, the test error exhibited an asymmetric U-shaped curve across training epochs, indicating limited generalizability. As the dataset size increased, training and test errors converged, with median mean absolute errors (MAE) of 0.041, 0.038, and 0.020 for the three dataset sizes, respectively. Bottom panels show MAE distributions across parameters, demonstrating progressively narrower error ranges as training datasets grow. **(C)** APs (V_m_), intracellular calcium transients (Cai), total ionic current (I_Total_), and six major individual ionic currents (I_Kr_, I_CaL_, I_Ks_, I_K1_, I_Na_, I_f_) were simulated from the predicted parameters of a single test cell using the deep learning network trained on 200,000 samples. Blue traces represent the original simulated data (input), and red traces show the model outputs. The close overlap confirms faithful reproduction of cell-specific electrophysiological behavior. See Figure S1 for a detailed comparison of the corresponding parameter values.

The deep learning network-generated parameters led to improved model waveforms (red) that duplicated ground truth simulated data (blue) for all outputs, with high fidelity in both amplitude and temporal features (**Figure 4C**). We observed overlap of AP morphology, Ca²⁺ transient dynamics, and the activation and inactivation profiles of each ion channel. These findings demonstrate that large, diverse training datasets are essential for high-fidelity parameter inference, enabling digital twins to accurately replicate both the quantitative dynamics and qualitative features of iPSC-CM electrophysiology. This scalability positions the framework for reliable application to experimental datasets where precise recovery of ionic properties is critical.

### Digital twins of human iPSC-CMs reproduce experimental electrophysiology and reveal intrinsic variability in drug response

While we have shown that the deep learning-based approach can create high-fidelity digital twins when trained and tested with in silico models of iPSC-CMs, the ultimate proving ground for the digital twin framework is application to real human iPSC-CM recordings. Therefore, we applied the optimized voltage-clamp protocol in its most demanding and key test to derive digital twins from real cells (**Figure 5**).

**Figure 5.**
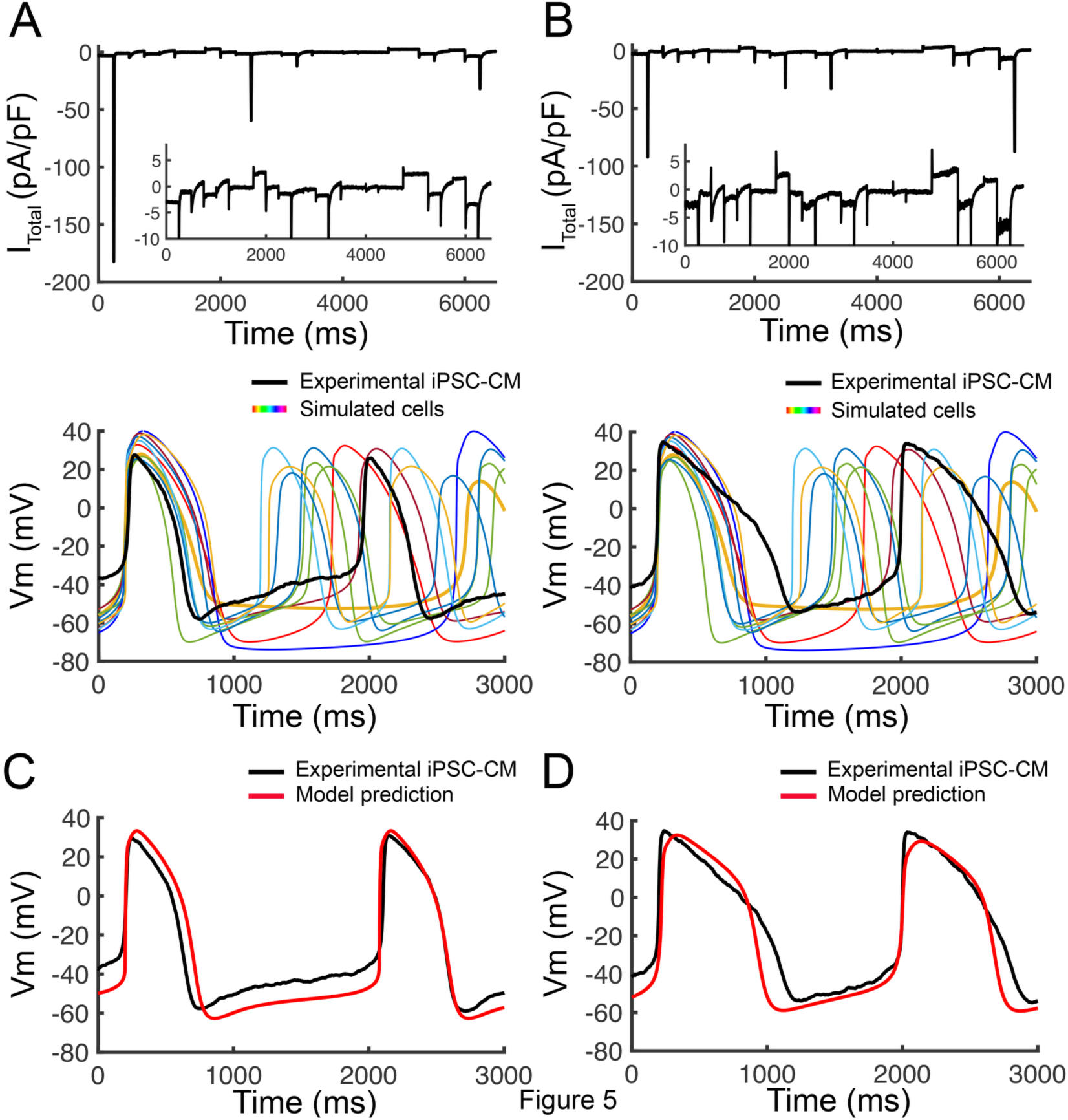
Digital twin generation and predictive modeling from real cells in iPSC-CM experimental recordings. (A, B, top) Whole-cell currents from two live iPSC-CMs in response to the optimized voltage-clamp protocol (Figure 3) were used as input to the trained deep learning network. Insets show expanded views of the responses. **(A, B, bottom)** Ten simulated APs from a synthetic population of 1,100,000 iPSC-CMs at room temperature were generated by introducing ±40% random variation to 52 biophysical model parameters from baseline, governing six major ionic currents: (I_Kr_, I_CaL_, I_Ks_, I_K1_, I_Na_, I_f_). Colored traces represent exemplar simulated cells from training data; overlaid black traces are experimentally recorded APs from two representative iPSC-CMs of the human iPSC cell line iPS-6-9-9T.B. (**C, D**) Digital twin models of the same experimental iPSC-CMs shown in panels A and B. Digital twins were created by extracting all 52 model parameters from the experimental whole-cell current using the deep learning inference pipeline. These parameters were used to instantiate cell-specific computational models, and AP simulations (red traces) were generated. The close overlay between the experimental traces (**black**) and digital twin predictions (**red**) demonstrates the success of the framework to accurately derive cell-specific digital twins from real cells.

The experimental cells in this study exhibited remarkably large variability in AP morphology, cycle length, and plateau characteristics, reflecting the heterogeneity typical of iPSC-CM preparations.

Capturing this breadth of behavior in the modeling pipeline required a correspondingly broad synthetic training dataset. To achieve this, we generated a large *in silico* population of 1,100,000 iPSC-CMs by applying ±40% random variation to 52 biophysical parameters governing six major ionic currents I_Kr_, I_CaL_, I_Na_, I_Ks,_ I_K1,_ and I_f_ relative to the baseline model. This expanded range ensured that extreme and rare phenotypes present in the experimental recordings were represented in the training space.

A second major consideration was temperature. While the iPSC-CM models were originally parameterized at 37 °C, the experimental data from iPSC-CM are commonly acquired at room temperature. To reconcile this mismatch, we systematically slowed all channel kinetics in the synthetic training population using a Q₁₀ factor of 3, a coarse but effective first approximation, to estimate the slower gating dynamics observed experimentally. This adjustment allowed the network to be trained on current waveforms that more closely matched the temporal characteristics of the experimental recordings.

Whole-cell currents recorded from two live iPSC-CMs (**Figure 5A and B, top**) in response to the protocol in Figure 3 served as the sole input to the trained deep learning network, which successfully inferred all 52 biophysical parameters associated with six major ionic currents, I_Kr_, I_CaL_, I_Na_, I_Ks,_ I_K1,_ and I_f_. These parameter sets were used to instantiate personalized digital twins for each experimental cell. Simulated APs (**A, B, bottom**) from the synthetic population at room temperature reflect the range of electrophysiological variability observed in experimental iPSC-CMs, indicating that the training data adequately capture the diversity of the experimental recordings.

When simulated, the digital twins reproduced the experimentally recorded APs with excellent agreement, capturing fine-scale features of depolarization rate, plateau amplitude and duration, and repolarization slope (**Figure 5C–D**), confirming that the pipeline reconstructs the full mechanistic basis of electrophysiology from a single experimental protocol.

To investigate how intrinsic variability among iPSC-CMs influences proarrhythmic susceptibility, we compared the responses of two representative model cells with distinct repolarization dynamics (**Figure 6A–B**). Under control conditions at 37 °C, both **Cell 1** and **Cell 2** generated stable APs with similar morphology. Exposure to the selective rapid delayed rectifier potassium current I_Kr_ blocker E-4031 (50 nM) markedly prolonged repolarization in both models, but only **Cell 1** exhibited early afterdepolarizations (EADs), whereas Cell 2 maintained stable repolarization. These findings highlight the strong impact of intrinsic cellular variability on proarrhythmic susceptibility and emphasize the value of digital twin models in capturing the heterogeneity of proarrhythmic responses to drug exposure across individual cells.

**Figure 6.**
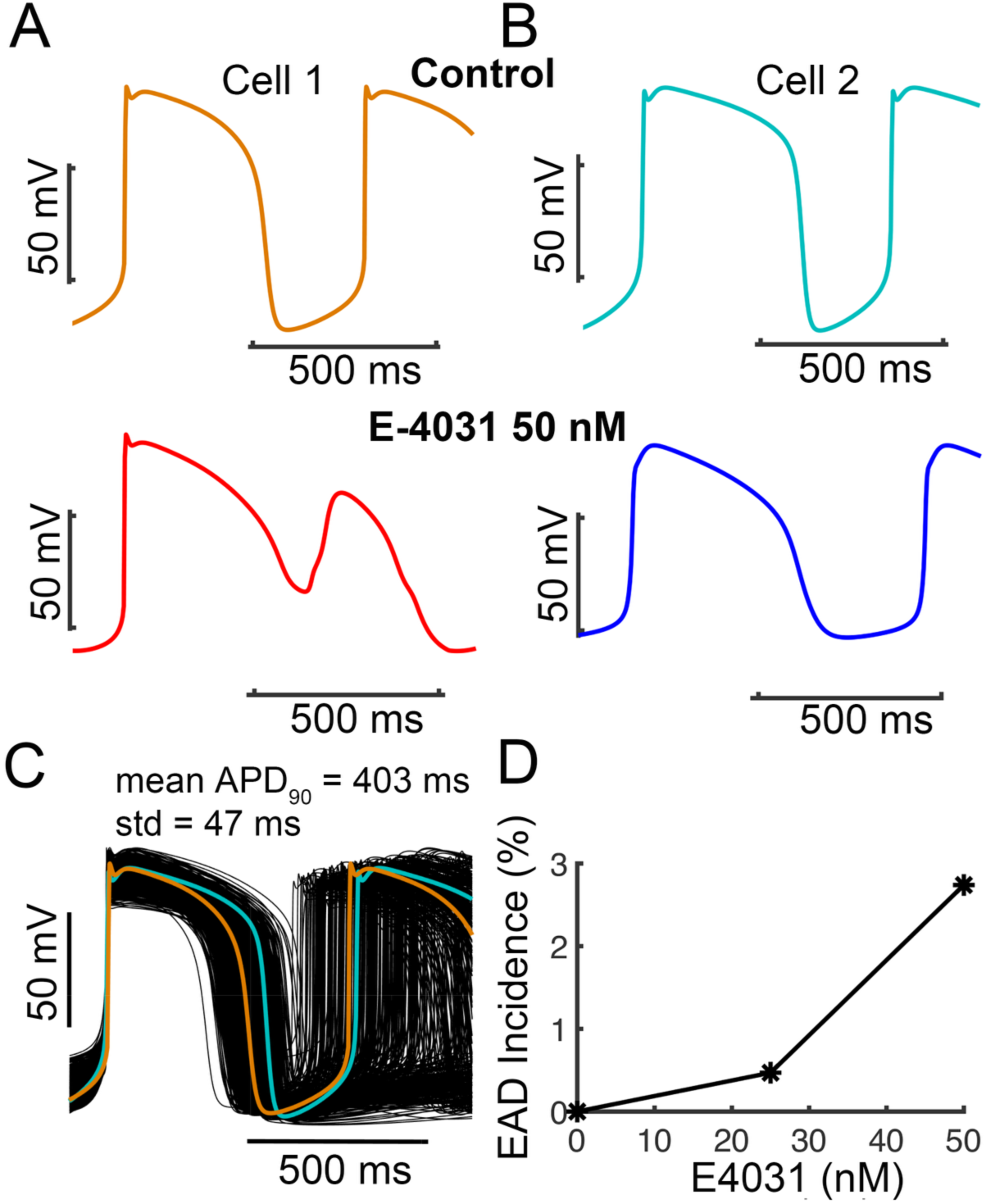
Cell-to-cell variability drives population-level responses to drug application in large synthetic iPSC-CM digital twins. (A) iPSC-CM model **Cell 1** (corresponding to Figure 5A). **(B)** iPSC-CM model **Cell 2** (corresponding to Figure 5B). In both panels, the **top traces** show control simulations at physiological temperature (37 °C). The **middle row** shows responses in the presence of E-4031 (50 nM). EADs occurred in **Cell 1** (red) at 50 nM, but not in **Cell 2** (blue). **(C)** We applied ±20% perturbations to all parameters governing six key ionic currents (I_K1_, I_Kr_, I_Ks_, I_CaL_, I_Na_, I_f_) in Cell 1 (orange) and Cell 2 (cyan) to generate a population of virtual cells (n = 4000), capturing the full spectrum of cell variability within a cell line. Overlaid membrane potential traces (black) show APs from Cell 1 and Cell 2. The population average action potential duration at 90% repolarization (APD_90_) was 403 ± 47 ms. **(D)** Incidence of EADs (%) in the virtual population as a function of E-4031 concentration.

To extend this analysis beyond single cells, we generated two *in silico* populations (n = 4000) derived from the network-predicted parameter sets of **Cell 1** and **Cell 2** (**Figure 6C**). Each population incorporated ±20% perturbations to all parameters governing six major ionic currents (I_Kr_, I_CaL_, I_Na_, I_Ks,_ I_K1,_ and I_f_), capturing the intrinsic cell-to-cell variability expected within a single human iPSC line. Under control conditions, both populations showed stable spontaneous firing with overlapping distributions of AP, reflecting baseline heterogeneity in electrophysiological properties.

The resulting population captured a broad distribution of repolarization phenotypes (mean APD_90_ = 403 ± 47 ms). Upon exposure to increasing concentrations of E-4031, the incidence of EADs increased in a dose-dependent manner, reaching approximately 3% of simulated cells at 50 nM (**Figure 6D**), based on pooled variants from both **Cell 1** and **Cell 2**. Although this represents a relatively small fraction of the population, it reflects the emergence of a susceptible subpopulation arising from intrinsic electrophysiological variability. These results demonstrate that modest cell-to-cell variability in ion-channel properties can substantially amplify heterogeneity in proarrhythmic responses within a population. The digital-twin framework, therefore, enables statistically robust quantification of arrhythmia risk by bridging single-cell electrophysiological diversity to population-level drug sensitivity.

## Discussion

This study demonstrates, for the first time, the development of digital twins of human iPSC-CMs directly from a single experimental recording that is readily generalizable to any excitable cell type. By combining an optimized voltage-clamp protocol, large-scale synthetic training populations, and deep learning–based parameter inference, we reconstructed the complete ionic parameter set underlying individual cell electrophysiology and reproduced AP and current waveforms with high fidelity. This study addresses a longstanding limitation in cardiac modeling: the inability to rapidly and accurately resolve the individualized ionic mechanisms that drive variability in both health and disease. The framework proved robust to two of the most challenging sources of variation in iPSC-CM studies, temperature differences and broad waveform diversity, highlighting the potential for a scalable and adaptable tool for basic research, preclinical safety testing, and precision medicine.

A broad range of AP morphologies and firing rates in our simulated iPSC-CM population were readily evoked with changes of ±40% across ionic parameters (**Figure 1**). This mirrors experimental reports of substantial heterogeneity in iPSC-CM electrophysiological properties, which may arise from differences in differentiation state, ion channel expression, and culture conditions^34–39^. Such diversity has important implications for both mechanistic interpretation and predictive modeling. It suggests that population-based approaches, rather than reliance on single-cell models, are essential for capturing the spectrum of cellular responses^40–43^.

The electrophysiological phenotype of the human iPSC-CMs in this study is represented using the Kernik–Clancy iPSC-CM model ^44^, which was developed to capture the range of AP morphologies and spontaneous behaviors observed experimentally within a single cell line ^37^. This population-based modeling framework incorporates variability in key ionic conductances to reproduce the spectrum of depolarization, repolarization, and automaticity characteristics commonly reported in human iPSC-CMs, while constraining parameter ranges to avoid non-physiological phenotypes.

Within this framework, we focused on six primary ionic currents that are well established as dominant determinants of AP shape and variability in iPSC-CMs. These currents govern critical features, including upstroke velocity, plateau dynamics, repolarization, and spontaneous activity, and their modulation has been shown to recapitulate experimentally observed cell-to-cell heterogeneity ^23,38,39,45–52^. Although many ionic currents are developmentally regulated and may vary across human iPSC-CM lines ^53^, this reduced set provides a parsimonious representation that captures the expected electrophysiological variability within a given cell line.

Importantly, the model is designed to be extensible. Additional currents or alternative parameterizations can be incorporated to reflect differences in maturation state, donor background, or differentiation protocol, enabling future studies to explicitly address variability across human iPSC-CM sources while maintaining consistency with the underlying framework.

Figure 2 highlights the key technology developed in this study, a deep learning-based workflow to recover the key ionic parameter set of a cell from a single voltage-clamp experiment. The synthetic training population of iPSC-CMs, generated by introducing random variation to 52 parameters governing six key ionic currents I_Kr_, I_CaL_, I_Na_, I_Ks,_ I_K1,_ and I_f_, ensured that the network was exposed to a broad spectrum of AP morphologies and whole-cell current profiles.

We developed a data-driven protocol design process guided by deep learning, as shown in Figure 3. In this approach, candidate voltage segments were iteratively evaluated to minimize parameter prediction error in synthetic populations, enabling the discovery of less variable, more targeted protocols. This optimization strategy effectively probed multiple ionic currents while reducing voltage variation, thereby improving voltage control and cell tolerance. The adaptive nature of the design process ensured that protocols were both information-rich and experimentally viable, providing a robust foundation for accurate parameter inference and digital twin construction from live iPSC-CM recordings. The iterative deep learning-guided approach to voltage-clamp protocol optimization offers clear advantages over traditional manual design strategies. Conventional protocols are typically based on heuristic knowledge of channel gating properties and are rarely tailored to maximize information for multi-parameter inference. Our adaptive framework directly couples experimental design with model performance metrics, using MSE as a quantitative feedback signal to iteratively refine the protocol. This ensures that each iteration selectively enriches the data with features most discriminative for parameter recovery, thereby accelerating convergence to an optimal waveform. Importantly, because the optimization is performed *in silico* using large-scale synthetic populations, the resulting protocol is both data-driven and generalizable, capturing a wide range of ionic phenotypes. Once developed, such a protocol can be implemented in experimental systems without modification, offering a powerful route to improve the precision and efficiency of digital twin construction in human cardiomyocytes.

The clear improvement in predictive accuracy and stability with increasing training dataset size, as shown in Figure 4, highlights a fundamental consideration for applying deep learning to biophysically detailed cardiac modeling. Larger datasets capture a wider range of morphological and kinetic variability in ionic currents, reducing the risk of model overfitting and enabling more reliable generalization to new experimental data. The largest dataset produced digital twins that match ground truth across APs, calcium dynamics, and individual currents, and indicates the value of comprehensive, diverse training data to ensure high-fidelity reproduction at the single-cell level.

The deep learning-guided protocol optimization framework described here lays the groundwork for the capstone result presented in Figure 5, where fully parameterized digital twins reproduced experimental iPSC-CM APs with excellent fidelity. By training on large-scale, information-rich synthetic datasets generated through optimized protocols, the platform achieves robustness to both temperature variation and broad morphological diversity in electrophysiological signals. This adaptability ensures that the same modeling pipeline can be rapidly retuned to diverse experimental conditions and phenotypes without sacrificing accuracy. As a result, the approach is not limited to *in vitro* modeling but is positioned for translational deployment, where personalized, physiology-grounded digital twins could guide patient-specific diagnostics, risk assessment, and therapy optimization. To move seamlessly from protocol design to clinically relevant predictive modeling represents a decisive step toward the integration of digital twins into the practice of precision cardiovascular medicine.

Figure 6 illustrates how intrinsic electrophysiological variability among iPSC-CMs can profoundly influence responses to pharmacological perturbation, even among cells derived from the same genetic background. Small differences in ionic conductance and gating kinetics were sufficient to determine whether a cell remained stable or developed EADs under identical drug exposure.

At the population level, the dose-dependent emergence of EADs provides a quantitative biomarker of proarrhythmic risk that cannot be captured by single deterministic simulations. Importantly, this emergent variability does not require extreme outlier phenotypes but arises naturally from multivariate perturbations around experimentally constrained parameter sets. This highlights the predictive value of large-scale digital twin populations for mechanistic safety pharmacology, enabling the translation of cell-level uncertainty into statistically robust risk metrics. Beyond E-4031, this framework can be generalized to compare patient-derived iPSC-CM lines or therapeutic perturbations, supporting the development of precision cardiotoxicity assessment pipelines that account for both genetic and stochastic sources of electrophysiological diversity.

The success of our parameter inference framework relies critically on the use of large, diverse synthetic datasets that could not be obtained through experimental recordings alone. Training the deep learning model required hundreds of thousands to millions of cellular signals. This is not possible to do experimentally as it would be prohibitively time-consuming, costly, and technically unfeasible to acquire high-fidelity voltage-clamp data at scale. In contrast, computationally generated data allow complete control over underlying parameters, precise labeling of ground truth values, and systematic exploration of parameter space well beyond what is directly observable in experiments. This approach not only accelerates model training but also ensures exposure to rare or extreme phenotypes and improves network generalization across diverse experimental cells.

In terms of scope, there are some limitations to our approach. The framework emphasizes six major ionic currents and a set of supporting mechanisms; however, it does not yet incorporate all ionic currents relevant to cardiac electrophysiology. Expanding the framework to include additional currents would necessitate introducing more parameters, which would increase both model complexity and the scale of training data required, although future iterations will undoubtedly address this limitation. Another limitation is the substantial computational resources needed to generate large synthetic datasets and train deep neural networks. Once training is complete, however, parameter inference is rapid and efficient, making the framework practical for downstream applications.

While developing and validating the digital twin framework *in silico* and *in vitro* using human iPSC-derived cardiomyocytes, the deep learning–guided protocol optimization framework can be extended to a wide range of excitable cell systems and experimental contexts. In principle, the same adaptive feedback loop could be deployed in patch-clamp or multi-electrode array recordings from any animal or human cell types, including neurons, smooth muscle cells, or engineered tissue constructs, where capturing rich ionic dynamics is essential for mechanistic modeling. Furthermore, by incorporating additional physiological variables such as calcium handling or mechano-electric feedback, the approach could evolve into a multi-modal protocol design tool capable of constraining both electrical and mechanical parameters in digital twin models.

This study examines a single iPSC-CM line across two independent differentiation batches, capturing cell-to-cell variability within each batch. While variability across lines, donors, and protocols was not directly evaluated here, the deep learning framework is structured to accommodate these additional sources of variation in future applications. The synthetic population was designed to span a broad range of physiologically plausible behaviors, providing a useful platform for probing potential response variability. While in this study we did not intend to represent a measured biological distribution, the framework can be readily extended for this purpose. The framework enables scalable prediction of population-level drug responses, and ongoing experimental efforts with larger and more diverse cell samples will further strengthen and refine the model predictions. Importantly, the approach is inherently extensible, and future studies incorporating replicate recordings across multiple cells, lines, and conditions will help deepen understanding of variability and expand its applicability.

We anticipate that near-future studies using the digital twin framework will enable high-throughput single-cell electrophysiological phenotyping for disease modeling, allowing investigators to map patient-derived cardiomyocyte function to individualized ionic mechanisms using a single experiment. The framework maintains accuracy across changes in temperature and accommodates wide variation in AP morphology, making it suitable for multicenter studies in which experimental conditions differ to improve reproducibility and enable harmonization of data. In preclinical safety assessment, temperature-independent digital twins could be used to evaluate the effects of new compounds at scale across a spectrum of cardiac phenotypes, identifying subgroups at increased risk. Looking further ahead, integration of this modeling platform with longitudinal patient data, including genetic information, imaging, and clinical outcomes, could give rise to living cardiac digital twins that evolve over time. Such models may ultimately support predictive diagnostics, optimization of therapy, and continuous monitoring in personalized healthcare.

## Methods

### Human iPSC-derived Ventricular Cardiomyocyte Differentiation and Culture

The human iPSC line iPS-6-9-9T.B was purchased from WiCell and cultured in StemMACS iPS-Brew XF medium (Miltenyi Biotec) on hESC-qualified Matrigel (Corning) at 37°C with 5% CO₂ and passaged at ∼70% confluency. Two independent batches of ventricular cardiomyocytes (VCMs) were differentiated in a 12-well plate by treating confluent human iPSCs with 6 µM CHIR99021 (Tocris) from day 0 to 1 (D0–D1) and 5 µM IWR-1 (Tocris) from day 2 to 4 (D2–D4) in RPMI 1640 supplemented with B27 without insulin (Thermo Fisher). From day 7 onward, human iPSC-derived VCMs were maintained in cardiomyocyte (CM) maintenance medium consisting of RPMI 1640 with B27 supplement with insulin. Cells were pooled from three wells of a 12-well plate and replated at approximately day 14 (D14) and maintained in CM maintenance medium supplemented with 2 µM CHIR99021 for an additional ∼7 days. Subsequently, cells were replated onto Matrigel-coated glass coverslips in CM maintenance medium for patch-clamp experiments. Patch-clamp recordings were performed on approximately day 21-25 (D21-25) post-differentiation. Recordings were obtained with cells sampled across 6 coverslips per batch under consistent experimental conditions.

### Experimental electrophysiology recordings

All electrophysiological recordings were performed at room temperature with an Axopatch 200A amplifier coupled to an Axon Digidata 1550B plus HumSilencer digitizer (Molecular Devices, San José, CA, USA). Cultured ventricular myocytes were placed on the stage of an inverted microscope (IX-71; Olympus Corporation, Tokyo, Japan) and continuously perfused with Tyrode’s solution. Patch pipettes were pulled from borosilicate glass capillaries (Sutter Instrument, Novato, CA, USA) and filled with the appropriate internal solution. The pipette solution contained (in mmol L⁻¹): 140 KCl, 5 NaCl, 10 EGTA, 5 Mg-ATP, and 10 HEPES, adjusted to pH 7.2 with KOH. Ionic currents were recorded in the whole-cell configuration of the patch-clamp technique at a sampling frequency of 10 kHz. After achieving the whole-cell configuration, the amplifier was switched to current-clamp mode for membrane potential measurements. The computationally optimized voltage-pulse protocols (see below) were used to efficiently probe membrane currents. For experimental validation datasets, any incomplete or failed recordings were excluded prior to analysis. Recordings were excluded if they exhibited no spontaneous firing, abnormally slow firing rates, or failed to capture a complete action potential waveform. These criteria were applied consistently across all recordings.

### Computationally optimized voltage-clamp protocol

A computationally optimized voltage-clamp protocol was designed to activate individual ionic currents while capturing their dynamic interactions (see Figure 3) sequentially and selectively. This protocol was applied to each synthetic cell to produce time-series data of total ionic current (I_Total_). Data were generated at a 0.1 ms time step and down-sampled to match the experimental acquisition rate. Following the derivation of the final optimized protocol, the time-series data of I_Total_ current was employed as input to the fully connected neural network, and the output layer contained 52 units corresponding to the model parameters, enabling the extraction of the model parameters.

### iPSC-CM electrophysiological model

A baseline iPSC-CM electrophysiological model was parameterized with 52 biophysical variables (see model equations in Supplementary Information) representing maximal conductances, gating kinetics, and calcium handling parameters for six major ionic currents (I_K1_, I_Kr_, I_Ks_, I_CaL_, I_Na_, I_f_). We selected ionic currents based on the Kernik-Clancy iPSC-CM model from our earlier research ^44^, which aims to simulate electrophysiological variability observed experimentally using a population-based approach. In this model, variability in key ionic currents was sufficient to generate the diverse action potential morphologies and spontaneous activities seen in iPSC-CMs, while avoiding non-physiological phenotypes. Using this approach, we concentrated on six primary ionic currents that critically influence action potential behavior and variability.

### Fully connected layers

Fully connected deep neural networks were implemented in TensorFlow using 26 hidden layers (1024 nodes per layer) with hyperbolic tangent (tanh) activation functions. Each layer performs a linear transformation followed by a nonlinear activation ^54,55^ :

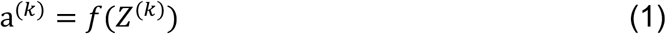

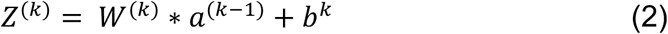

where 𝑎^(*k*)^ is the activation vector at layer 𝑘, 𝑊^(*k*)^ and 𝑏^(*k*)^ are the weight matrix and bias vector, respectively, and 𝑓 denotes the nonlinear activation function.

The network input is the time series of 𝐼*_Total_*(𝑡), obtained from the voltage-clamp protocol. The network outputs a vector of predicted biophysical parameters, **p̂** ∈ ℝ^+,^, representing ionic conductances and gating kinetics.

### Training and loss function

Networks were trained using mean squared error (MSE) loss and the Adam optimizer (1 × 10⁻⁴). MSE was used as the function evaluation metric (Eq. 3) for predicted model parameters ^56,57^.

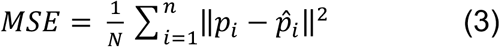

where *N* is the number of samples, *p_i_* is the ground-truth parameter vector, and *p̂_i_* is the corresponding predicted parameters (the network output). To illustrate how dataset size influences predictive accuracy, a narrower perturbation range of ±20% was used to measure variability around the baseline parameters (Figure 4). We independently sampled 1,000, 10,000, and 200,000 cells at 310 Kelvin (37 °C) to assess how dataset size impacts prediction performance.

For a comprehensive exploration of model variability, a wider ±40% perturbation range was chosen for the 52 parameters (Figure 5). Additionally, all channel kinetics in the synthetic training set were slowed using a Q₁₀ factor of 3 to simulate conditions at 298 K (room temperature). The 1.1M-cell dataset size was chosen to ensure broad exploration of the parameter space. This approach helps capture a broad spectrum of experimental differences while ensuring the ionic current properties and action potential behavior stay within realistic limits. Since the full model involved changing many parameters at once, we used a wider range to ensure a diverse enough population, but not so large that the solutions would become mostly non-physiological. Due to the broad parameter variation (±40%), some simulated cells exhibited non-physiological behavior, including non-excitable and non-repolarizing cells. These cells were excluded from the dataset, and only models demonstrating physiologically relevant electrophysiological activity and numerically stable behavior were retained for analysis.

We split the data into training, validation, and test sets using a 70:10:20 ratio, and implemented the network architecture using Keras ^58^. The dataset was also divided into 1% sample batches, requiring 70 iterations per epoch. As we monitored the evaluation metrics using the MSE, we continually optimized the voltage clamp protocol to improve the error output and, therefore, get continuously closer to an optimal data generation algorithm for parameter extraction. To prevent overfitting, we computed evaluation metrics using validation data at each training iteration and compared them to values from the training data. When model performance on the training data began to degrade relative to the validation dataset, tuning of the network hyperparameters stopped. Model accuracy was assessed using median mean absolute error (MAE) across parameters. This approach allowed us to extract key ion channel parameters from whole-cell currents, ensuring that the reconstructed ionic current and AP traces closely mirror the physiological behavior of individual cardiac cells.

### Population generation and drug testing

We generated a population by applying ± 20% perturbations to all parameters associated with six major ionic currents (I_K1_, I_Kr_, I_Ks_, I_CaL_, I_Na_, I_f_) around the network-predicted parameter sets for Cell 1 and Cell 2, both originating from the human iPSC lines iPS 6-9-9T.B. This procedure produced a 4000-cell population capturing within-line variability. For the EAD population, we used a narrower perturbation range (±20%) to focus on the parameter space near the baseline model. This allowed us to examine EAD and highlight parameter combinations that promote EAD formation, which only required (±20%) perturbation. The cell population captures intrinsic cell-to-cell variability within a shared genetic background. Simulations were performed under control conditions and in the presence of E-4031 concentrations using a Hill-type hERG block model.

### Simulation of I_Kr_ Blockade

To simulate the inhibitory effects of E-4031 on I_Kr_ current, we decreased the peak conductance, G_Kr_ of each of these independent channels in a concentration-dependent fashion using a concentration response relationship with a Hill coefficient of 1 (n = 1) as follows:

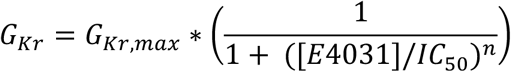

where G_Kr,max_ is the nominal conductance value obtained from each ventricular myocyte model, and the IC_50_ (7 nM) ^44,59^ is the concentration of drug that produces a 50% inhibition of the targeted transmembrane current or intracellular organelle.

Simulation and analysis code was written in C++, Python, TensorFlow and MATLAB R2018a (The MathWorks, Natick, MA, USA). The code was executed on two systems: a BIZON ZX5500 workstation with a 2.7 GHz 64-core AMD Threadripper Pro processor, an NVIDIA H100 GPU, and Python version 3.12, TensorFlow version 2.19; and a Mercury GPU408 server with a 2.55 GHz 192-core AMD EPYC 9684 processor, four NVIDIA H100 GPUs, and Python version 3.10, TensorFlow version 2.18. Compilation was performed using GCC version 10.5. Numerical results were visualized in MATLAB R2018a.

## Data availability

Experimental dataset generated for this study is available at https://github.com/ClancyLabUCD/Digital-Twin-for-the-Win-Personalized-Cardiac-Electrophysiology).

## Code availability

Due to the large size of the synthetic datasets (∼1.1M cells), the raw data are not hosted online. However, a representative dataset of 100 cells is provided as an example, and the complete codebase for dataset generation, along with parameter distributions and simulation protocols, is publicly available at GitHub (https://github.com/ClancyLabUCD/Digital-Twin-for-the-Win-Personalized-Cardiac-Electrophysiology). This allows full regeneration of the training and validation datasets as described in the study.

## Sources of funding

This work was supported in part by the National Institutes of Health under grants R01HL128537, OT2OD026580, and R01HL17400 (to C. E. Clancy and L. F. Santana), and R01HL159492 (to D. K. Lieu). G. H-Hernandez was supported by the UC Davis Chancellor’s Postdoctoral Fellowship. Additional support from The UC Davis Center for Precision Medicine and Data Sciences was provided to C. E. Clancy, P.-C. Yang and computing facilities.

## Disclosures

D.K. Lieu serves as a scientific consultant for Novoheart, Ltd.

## Supplementary Information

### iPSC-CM electrophysiological model gating kinetics parameters

**Figure S1.**
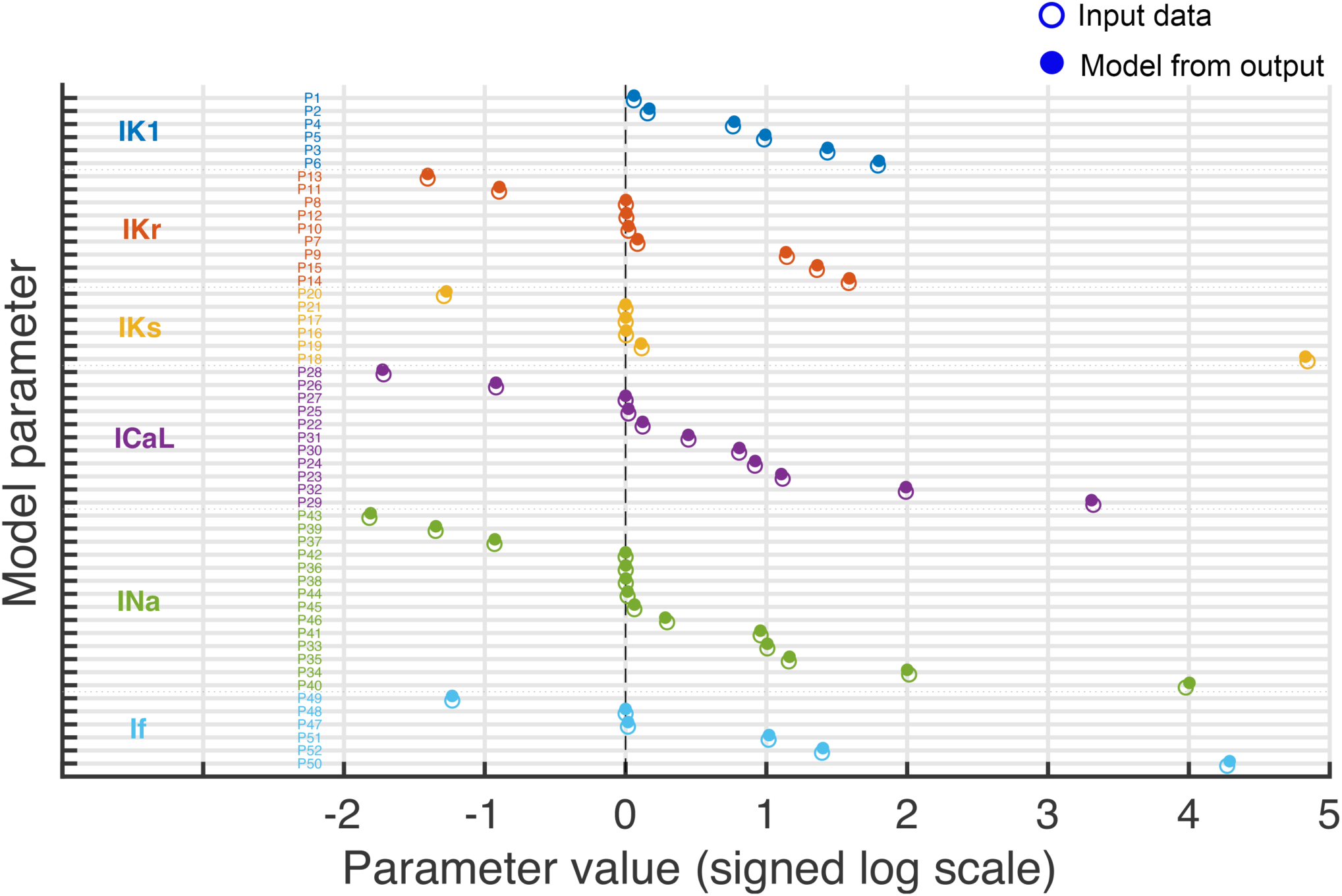
Comparison of model parameter values before and after perturbation. Each parameter (see Table 1 below) is represented by two markers: open circles denote baseline values and filled circles denote updated values. Parameters are grouped by ionic current (IK1, IKr, IKs, ICaL, INa, If), with colors indicating group membership. Within each group, parameters are ordered by the transformed value of the updated condition. Values are displayed on a signed logarithmic scale (sign(x)·log₁₀(1+|x|)) to accommodate both positive and negative values across a wide dynamic range. The vertical dashed line indicates zero. Horizontal dotted lines separate parameter groups.

**Table 1:**
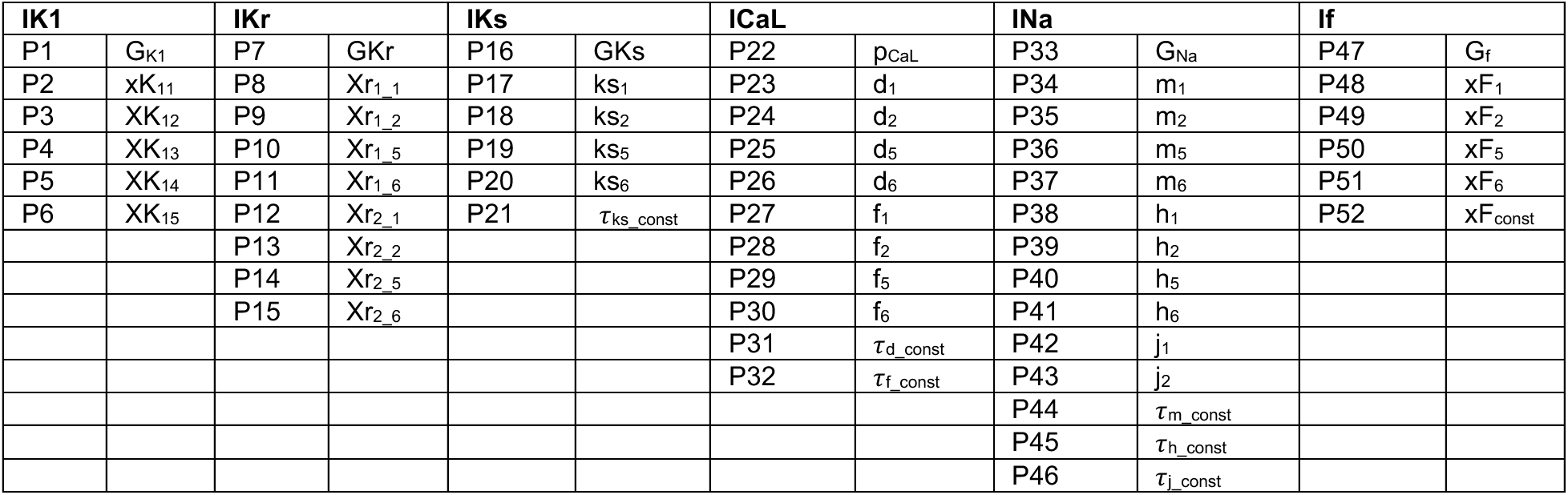
Definitions and correspondence of model parameters (P1–P52) with the variables used in the mathematical equations below.

### iPSC-CM electrophysiological model equations

The 52 biophysical variables are highlighted in red in the equations below.

#### Inward rectifier potassium current (I_K1_)

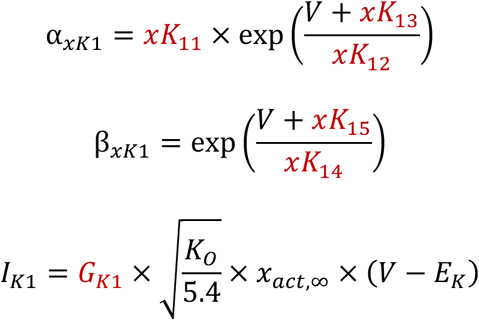

#### Rapid delayed rectifier potassium current (I_Kr_)

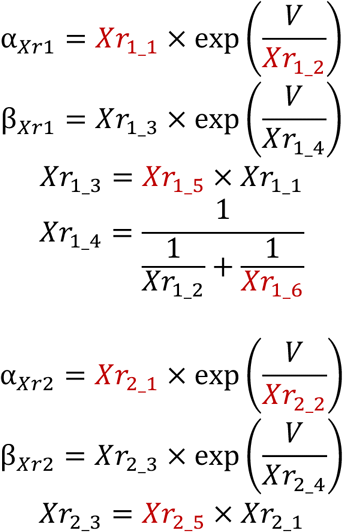

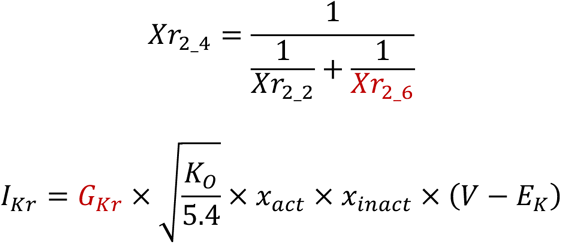

#### Slow delayed rectifier potassium current (I_Ks_)

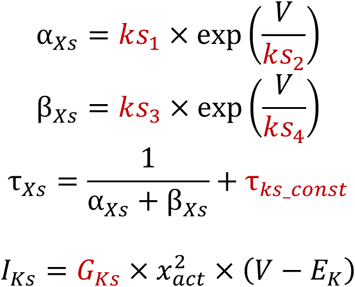

#### L-type Ca^2+^ current (I_CaL_)

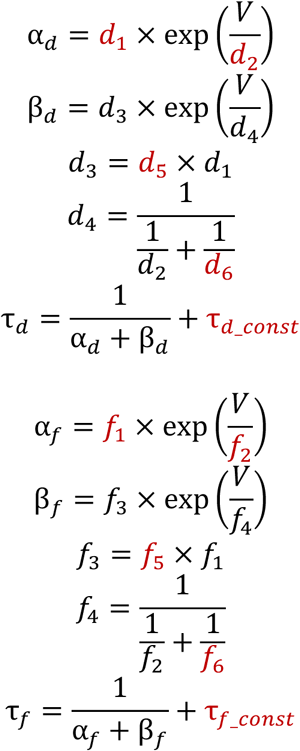

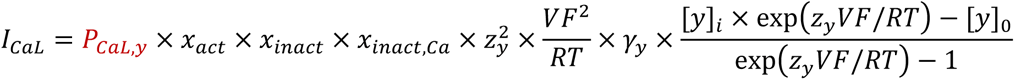

Where *y* is Ca^2+^, Na^+^, or K^+^, 𝑃_B)C_ is the permeability ion *y*, *R* is the gas constant, *z*_y_ is the valence of ion *y*, and γ_y_ is to activity coefficient for ion *y*.

#### Sodium Current (I_Na_)

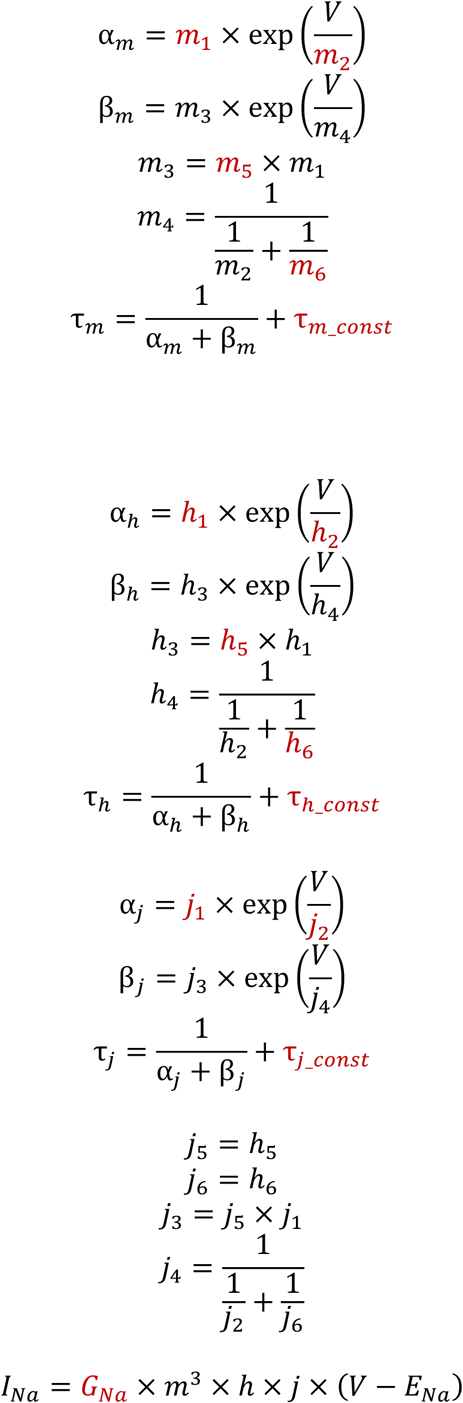

#### Funny/HCN current (I_f_)

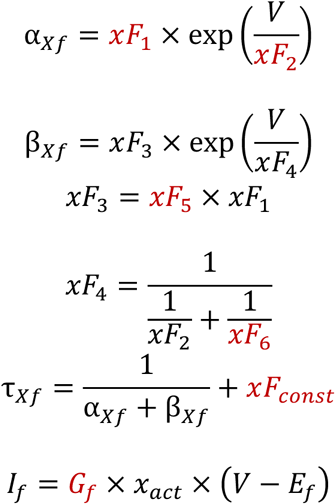

For detailed model equations and parameters, please refer to the Kernik-Clancy model^1^.

## Notes

### Summary of Updates

The revised version now includes the supplementary material within the main article file.

