## Supplementary Information for "Large-scale synthetic data enable digital twins of human excitable cells"

#### iPSC-CM electrophysiological model gating kinetics parameters

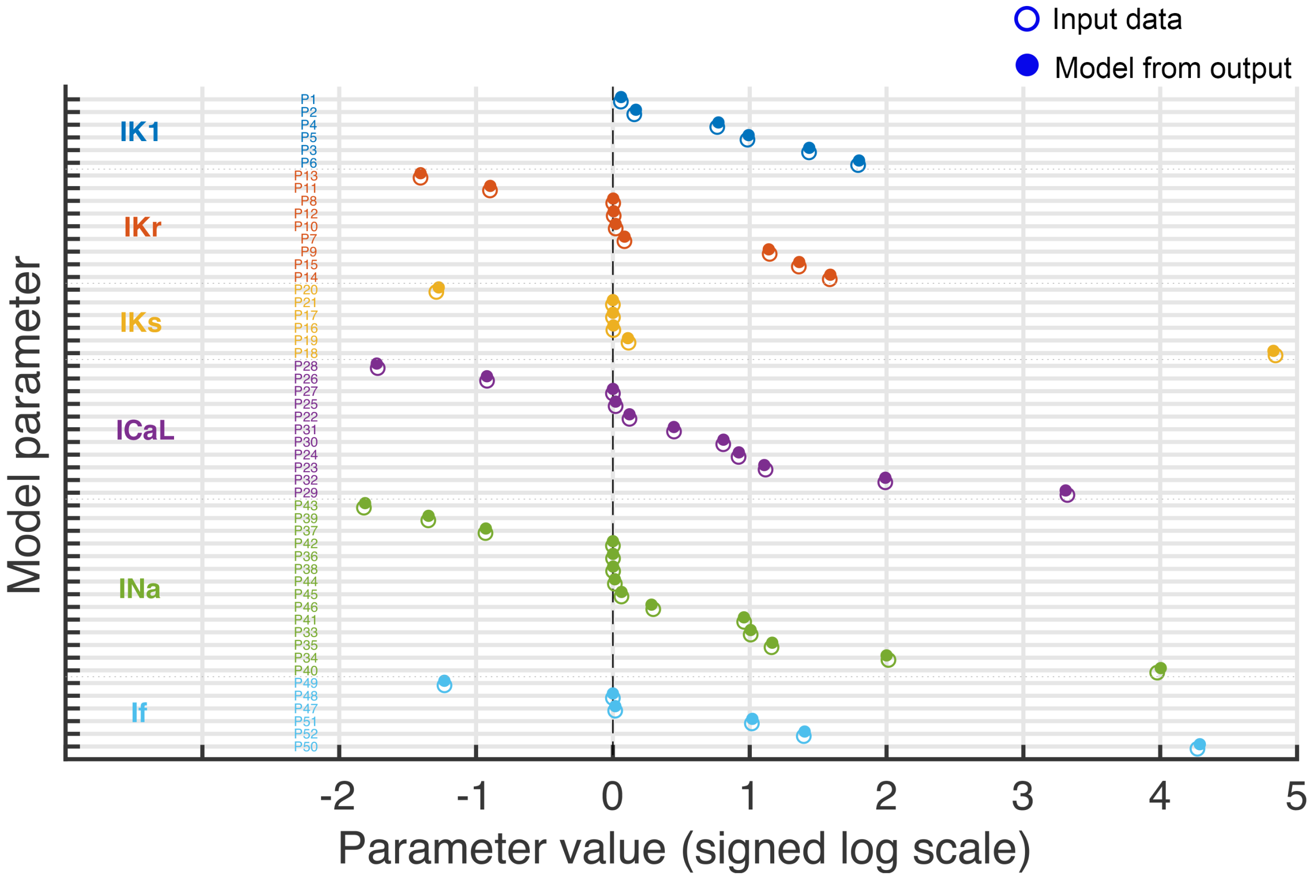


**Figure S1. Comparison of model parameter values before and after perturbation**. Each parameter (see Table 1 below) is represented by two markers: open circles denote baseline values and filled circles denote updated values. Parameters are grouped by ionic

current (IK1, IKr, IKs, ICaL, INa, If), with colors indicating group membership. Within each group, parameters are ordered by the transformed value of the updated condition. Values are displayed on a signed logarithmic scale ($\text{sign}(x)\cdot\log_{10}(1+|x|)$) to accommodate both positive and negative values across a wide dynamic range. The vertical dashed line indicates zero. Horizontal dotted lines separate parameter groups.

Table 1: Definitions and correspondence of model parameters (P1–P52) with the variables used in the mathematical equations below.

| **IK1** | | **IKr** | | **IKs** | | **ICaL** | | **INa** | | **If** | |
|---|---|---|---|---|---|---|---|---|---|---|---|
| P1 | $G_{K1}$ | P7 | GKr | P16 | GKs | P22 | $p_{CaL}$ | P33 | $G_{Na}$ | P47 | $G_f$ |
| P2 | $xK_{11}$ | P8 | $Xr_{1\_1}$ | P17 | $ks_1$ | P23 | $d_1$ | P34 | $m_1$ | P48 | $xF_1$ |
| P3 | $XK_{12}$ | P9 | $Xr_{1\_2}$ | P18 | $ks_2$ | P24 | $d_2$ | P35 | $m_2$ | P49 | $xF_2$ |
| P4 | $XK_{13}$ | P10 | $Xr_{1\_5}$ | P19 | $ks_5$ | P25 | $d_5$ | P36 | $m_5$ | P50 | $xF_5$ |
| P5 | $XK_{14}$ | P11 | $Xr_{1\_6}$ | P20 | $ks_6$ | P26 | $d_6$ | P37 | $m_6$ | P51 | $xF_6$ |
| P6 | $XK_{15}$ | P12 | $Xr_{2\_1}$ | P21 | $\tau_{ks\_const}$ | P27 | $f_1$ | P38 | $h_1$ | P52 | $xF_{const}$ |
| | | P13 | $Xr_{2\_2}$ | | | P28 | $f_2$ | P39 | $h_2$ | | |
| | | P14 | $Xr_{2\_5}$ | | | P29 | $f_5$ | P40 | $h_5$ | | |
| | | P15 | $Xr_{2\_6}$ | | | P30 | $f_6$ | P41 | $h_6$ | | |
| | | | | | | P31 | $\tau_{d\_const}$ | P42 | $j_1$ | | |
| | | | | | | P32 | $\tau_{f\_const}$ | P43 | $j_2$ | | |
| | | | | | | | | P44 | $\tau_{m\_const}$ | | |
| | | | | | | | | P45 | $\tau_{h\_const}$ | | |
| | | | | | | | | P46 | $\tau_{j\_const}$ | | |

### iPSC-CM electrophysiological model equations

The 52 biophysical variables are highlighted in red in the equations below.

*Inward rectifier potassium current ($I_{K1}$)*

$$\alpha_{xK1} = xK_{11} \times \exp\left(\frac{V + xK_{13}}{xK_{12}}\right)$$

$$\beta_{xK1} = \exp\left(\frac{V + xK_{15}}{xK_{14}}\right)$$

$$I_{K1} = G_{K1} \times \sqrt{\frac{K_O}{5.4}} \times x_{act,\infty} \times (V - E_K)$$

*Rapid delayed rectifier potassium current ($I_{Kr}$)*

$$\alpha_{Xr1} = Xr_{1\_1} \times \exp\left(\frac{V}{Xr_{1\_2}}\right)$$

$$\beta_{Xr1} = Xr_{1\_3} \times \exp\left(\frac{V}{Xr_{1\_4}}\right)$$

$$Xr_{1\_3} = Xr_{1\_5} \times Xr_{1\_1}$$

$$Xr_{1\_4} = \frac{1}{\frac{1}{Xr_{1\_2}} + \frac{1}{Xr_{1\_6}}}$$

$$\alpha_{Xr2} = Xr_{2\_1} \times \exp\left(\frac{V}{Xr_{2\_2}}\right)$$

$$\beta_{Xr2} = Xr_{2\_3} \times \exp\left(\frac{V}{Xr_{2\_4}}\right)$$

$$Xr_{2\_3} = Xr_{2\_5} \times Xr_{2\_1}$$

$$Xr_{2\_4} = \frac{1}{\frac{1}{Xr_{2\_2}} + \frac{1}{Xr_{2\_6}}}$$

$$I_{Kr} = G_{Kr} \times \sqrt{\frac{K_O}{5.4}} \times x_{act} \times x_{inact} \times (V - E_K)$$

*Slow delayed rectifier potassium current ($I_{Ks}$)*

$$\alpha_{Xs} = ks_1 \times \exp\left(\frac{V}{ks_2}\right)$$

$$\beta_{Xs} = ks_3 \times \exp\left(\frac{V}{ks_4}\right)$$

$$\tau_{Xs} = \frac{1}{\alpha_{Xs} + \beta_{Xs}} + \tau_{ks\_const}$$

$$I_{Ks} = G_{Ks} \times x_{act}^2 \times (V - E_K)$$

*L-type $Ca^{2+}$ current ($I_{CaL}$)*

$$\alpha_d = d_1 \times \exp\left(\frac{V}{d_2}\right)$$

$$\beta_d = d_3 \times \exp\left(\frac{V}{d_4}\right)$$

$$d_3 = d_5 \times d_1$$

$$d_4 = \frac{1}{\frac{1}{d_2} + \frac{1}{d_6}}$$

$$\tau_d = \frac{1}{\alpha_d + \beta_d} + \tau_{d\_const}$$

$$\alpha_f = f_1 \times \exp\left(\frac{V}{f_2}\right)$$

$$\beta_f = f_3 \times \exp\left(\frac{V}{f_4}\right)$$

$$f_3 = f_5 \times f_1$$

$$f_4 = \frac{1}{\frac{1}{f_2} + \frac{1}{f_6}}$$

$$\tau_f = \frac{1}{\alpha_f + \beta_f} + \tau_{f\_const}$$

$$I_{CaL} = P_{CaL,y} \times x_{act} \times x_{inact} \times x_{inact,Ca} \times z_y^2 \times \frac{VF^2}{RT} \times \gamma_y \times \frac{[y]_i \times \exp(z_y VF/RT) - [y]_0}{\exp(z_y VF/RT) - 1}$$

Where $y$ is $Ca^{2+}$, $Na^+$, or $K^+$, $P_{CaL}$ is the permeability ion $y$, $R$ is the gas constant, $z_y$ is the valence of ion $y$, and $\gamma_y$ is to activity coefficient for ion $y$.

*Sodium Current ($I_{Na}$)*

$$\alpha_m = m_1 \times \exp\left(\frac{V}{m_2}\right)$$

$$\beta_m = m_3 \times \exp\left(\frac{V}{m_4}\right)$$

$$m_3 = m_5 \times m_1$$

$$m_4 = \frac{1}{\frac{1}{m_2} + \frac{1}{m_6}}$$

$$\tau_m = \frac{1}{\alpha_m + \beta_m} + \tau_{m\_const}$$

$$\alpha_h = h_1 \times \exp\left(\frac{V}{h_2}\right)$$

$$\beta_h = h_3 \times \exp\left(\frac{V}{h_4}\right)$$

$$h_3 = h_5 \times h_1$$

$$h_4 = \frac{1}{\frac{1}{h_2} + \frac{1}{h_6}}$$

$$\tau_h = \frac{1}{\alpha_h + \beta_h} + \tau_{h\_const}$$

$$\alpha_j = j_1 \times \exp\left(\frac{V}{j_2}\right)$$

$$\beta_j = j_3 \times \exp\left(\frac{V}{j_4}\right)$$

$$\tau_j = \frac{1}{\alpha_j + \beta_j} + \tau_{j\_const}$$

$$j_5 = h_5$$

$$j_6 = h_6$$

$$j_3 = j_5 \times j_1$$

$$j_4 = \frac{1}{\frac{1}{j_2} + \frac{1}{j_6}}$$

$$I_{Na} = G_{Na} \times m^3 \times h \times j \times (V - E_{Na})$$

*Funny/HCN current ($I_f$)*

$$\alpha_{Xf} = xF_1 \times \exp\left(\frac{V}{xF_2}\right)$$

$$\beta_{Xf} = xF_3 \times \exp\left(\frac{V}{xF_4}\right)$$

$$xF_3 = xF_5 \times xF_1$$

$$xF_4 = \frac{1}{\frac{1}{xF_2} + \frac{1}{xF_6}}$$

$$\tau_{Xf} = \frac{1}{\alpha_{Xf} + \beta_{Xf}} + xF_{const}$$

$$I_f = G_f \times x_{act} \times \left(V - E_f\right)$$

For detailed model equations and parameters, please refer to the Kernik-Clancy model[1].
